# Preclinical evaluation of [^11^C]CHDI-009R for quantification of mutant huntingtin aggregates

**DOI:** 10.1101/2025.06.17.660088

**Authors:** Franziska Zajicek, Liesbeth Everix, Annemie Van Eetveldt, Jeroen Verhaeghe, Stef De Lombaerde, Alan Miranda, Jordy Akkermans, Celia Dominguez, Vinod Khetarpal, Jonathan Bard, Longbin Liu, Steven Staelens, Daniele Bertoglio

## Abstract

Huntington’s disease (HD) is a neurodegenerative disorder caused by an expanded trinucleotide repeat in the huntingtin gene (*HTT*) that subsequently leads to aggregation of the mutant huntingtin (mHTT) protein. Thus, lowering mHTT is a key therapeutic approach used by several candidate therapeutics currently under investigation. Visualization of the efficiency of these therapeutics through *in vivo* mHTT quantification rises in importance. For positron emission tomography (PET) imaging of mHTT aggregates, it is critical to characterize the *in vivo* kinetic profile of newly identified mHTT binders to assess their translational application. Here, we report the evaluation of [^11^C]CHDI-009R, a PET imaging radioligand with higher affinity and selectivity for mHTT aggregates than previously reported radioligands, in the heterozygous (HET) zQ175DN mouse model of HD and wild-type (WT) littermates at 9 and 3 months of age.

[^11^C]CHDI-009R displayed high stability in plasma and brain, which was reflected in brain kinetics as demonstrated by rapid uptake followed by relatively slow elimination. Kinetic modeling and volume of distribution *V*_T (IDIF)_ indicated the radioligand’s ability to quantify mHTT aggregation at 9 months of age with clear genotype differentiation (*p*<0.0001). [^11^C]CHDI-009R showed an excellent test-retest reliability in 9-month-old mice (intraclass correlation coefficient: 0.62 - 0.79). A phenotypic difference in mHTT aggregates was also observed in 3-month-old mice in several brain structures (*p*<0.05) and was confirmed with [^3^H]CHDI-009R autoradiography.

Overall, this study suggests [^11^C]CHDI-009R is a promising radioligand for the detection of cerebral mHTT aggregates in a mouse model of HD and supports its advance to clinical evaluation.

## Introduction

Huntington’s disease (HD) is a progressive neurodegenerative disorder caused by an expanded trinucleotide (CAG) repeat in exon 1 of the huntingtin (*HTT*) gene. The transcription and translation of mutant *HTT* leads to the aggregation of mutant huntingtin (mHTT) protein [1, 2], which, alongside mHTT fragments, is involved in cellular dysfunction and loss of neurons [3, 4]. With several disease-modifying therapeutic interventions aimed to lower mHTT expression under clinical investigation [5–7] (e.g. ClinicalTrials.gov identifier (ID): NCT06024265; NCT05032196), there is an increasing interest in accurately quantifying their ameliorative effect. Currently, the estimate of the central nervous system (CNS) target engagement of HTT-lowering therapeutics is indirect, relying on extrapolations of soluble HTT levels in non-brain compartments, such as cerebrospinal fluid, both in the preclinical [8] and clinical setting [9] (www.clinicaltrials.gov, identifier NCT02519036). Still, fluid biomarkers lack brain region-specific information that positron emission tomography (PET) imaging of mHTT aggregates could provide. Thus, PET imaging holds great potential as a complementary tool to assess brain region-specific target engagement, as previously demonstrated preclinically [10]. Thus, enabling accurate spatial and temporal evaluation of efficacy with potential therapeutic interventions.

We recently described a number of novel radiotracers for mHTT aggregate imaging in preclinical imaging studies, namely [^11^C]CHDI-626 [11–13], [^11^C]CHDI-180R [10, 12, 14, 15] and [^18^F]CHDI-650 [16, 17] with varying levels of applicability in mice. Additionally, [^11^C]CHDI-626 [18] was advanced to the clinic but later discontinued due to an undesirable brain uptake and decay profile. On the other hand, [^11^C]CHDI-180R detected a significant difference between people with HD (PwHD) and healthy controls; however, it also displayed high intersubject variability [19]. Thus, a more suited clinical candidate would be preferred, and the identification of novel radioligands targeting mHTT aggregates with superior PET radiotracer properties remains a priority. Especially regarding clinical importance, enhanced sensitivity of mHTT aggregate detection in 3-month-old zQ175DN heterozygous (HET) mice is desired, as mHTT aggregation is expected to be lower in abundance in humans than in mouse models. Therefore, we report on the investigation of a novel radioligand, [^11^C]CHDI-009R, that has been shown to have an almost 3-fold higher maximum binding capacity than CHDI-180R based on autoradiography studies in homozygous zQ175DN brains [20]. Here, we characterize the *in vivo* performance of this tracer in wildtype (WT) and HET zQ175DN knock-in mice through radiometabolite analysis, test-retest variability estimation, quantitative phenotypic comparison at two different ages, and post-mortem autoradiography.

## Methods

### Animals

A total of 63 nine-month-old male HET (*n*=33) and WT (*n*=30) as well as 44 three-month-old male HET (*n*=24) and WT (*n*=20) zQ175DN mice (B6.129S1-*Htt^tm1.1Mfc^*/190ChdiJ JAX stock #029928), were obtained from Jackson Laboratories (Bar Harbour, Maine, USA) [21, 22]. Due to sporadic congenital portosystemic shunt occurring in C57BL/6J mice [23], all animals were screened at Jackson Laboratories before shipment, and hence, all animals used in this study were shunt-free. Study design, group size, body weight, and age of the animals are reported in Fig. 1 and Tables S1-S3. Animals were single-housed at 9 months of age and group-housed at 3 months of age in ventilated cages under a 12 h light/dark cycle. While group-housing is generally considered to achieve better social well-being for mice, minimising aggression and potential injuries caused by dominant-submissive behaviour was prioritised for animals undergoing several imaging sessions. The animals were kept in a temperature- and humidity-controlled environment with food and water *ad libitum*. The animals were given at least one week for acclimatization after arrival before the start of any procedure. Experiments followed the European Committee (decree 2010/63/CEE) and were approved by the Ethical Committee for Animal Testing (ECD 2019-39) at the University of Antwerp (Belgium).

**Fig. 1.**
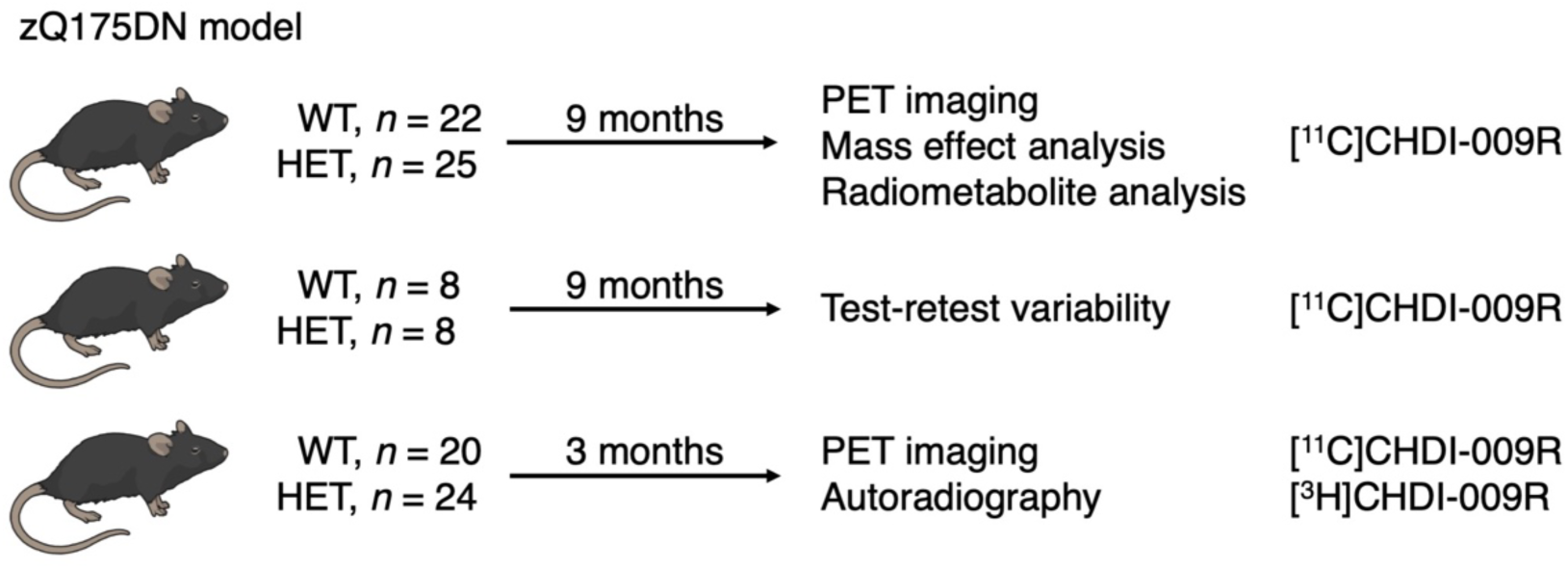
Overview of the experimental design. Number and age of each experimental group for the different experimental setups. HET = heterozygous; n = number; WT = wildtype

### Radioligand synthesis

[^11^C]CHDI-009R was prepared using an automated synthesis module (Carbonsynthon I, Comecer, The Netherlands). The radioligand was prepared via single-step carbon-11 labeling at the pyridazinone-NH site, starting with a solution of the precursor (0.3 mg) in DMF (0.5 ml), which was reacted with excess of [^11^C]CH_3_OTf in the presence of NaOH (1 M, 5 µl) for 3 min at room temperature. To terminate the reaction and to ensure good retention of the compound of interest on the semi-preparative HPLC column, acetonitrile/water (35:65, v/v, 2 ml) was introduced to the reaction mixture, and the resulting crude product was purified on HPLC using an ACE 5 C18-HL 250×10 mm column (ACE, UK), eluted with a mixture of ammonium formate (0.1 M) in acetonitrile (35/65, v/v) as mobile phase, at a flow rate of 6 ml/min. The collected fraction was diluted with water (20 ml) before being loaded on a 1 cc 10 mg Oasis HLB cartridge (Waters, MA, USA). The cartridge was washed with water for injection (3 ml) and eluted through a sterile filter with ethanol (0.5 ml) and saline (4.5 ml) to afford the final product solution.

The radiochemical purity of the produced [^11^C]CHDI-009R was determined using a Kinetex EVO C18, 5µm, 150×4.6 mm (Phenomenex, USA) HPLC column, with a solution of sodium acetate (NaOAc) (0.05 M, pH 5.5) in acetonitrile/ (30:70, v/v) as mobile phase, at a flow of 1 ml/min, with UV absorbance set at 289 nm. Based on 37 productions, radiochemical purity was greater than 99%, a molar activity (A_m_) of 93.59±27.61 GBq/µmol, and a radiochemical yield of 7.83±0.03% was achieved.

### Radiometabolite analysis

Evaluation of *in vivo* plasma and brain radiometabolite profiles was performed at 5, 20, and 40 min post-injection (p.i.). Mice were injected via the lateral tail vein with [^11^C]CHDI-009R (WT: 3.78±0.87 MBq; HET: 3.75±0.62 MBq in 200 μl), and blood was withdrawn via cardiac puncture at designated times, and brains were immediately excised. The homogenization and extraction were conducted as previously described [10, 17]. The extraction efficiency from plasma samples (percentage of recovery of radioactivity) was 98.4±2.6%, whereas in brain samples, an extraction efficiency of 100.6±4.7% was achieved. Plasma or brain supernatant was loaded onto a pre-conditioned reverse-phase (RP)-HPLC system (Luna C18(2), 5 μm HPLC column (250×4.6 mm) + Phenomenex security guard pre-column) and eluted with NaOAc 0.05M pH 5.5 and acetonitrile: (87:13 for 10 min v/v) buffer at a flow rate of 1 ml/min. RP-HPLC fractions were collected at 0.5 min intervals for 10 min. Pure tracer (5µCi) was loaded onto the same HPLC setup (Fig. S1). Control experiments confirmed that no apparent degradation of the radiotracer occurred during procedural work-up in both plasma (99.91±0.00% of intact tracer) and brain (92.14±0.07% of intact tracer).

### Dynamic PET imaging acquisition

Dynamic microPET/Computed tomography (CT) images were acquired on Siemens Inveon PET-CT scanners (Siemens Preclinical Solution, Knoxville, USA). Animal preparation was performed as previously described [10, 24]. Animal and dosing information for both WT and HET mice of all paradigms are given in the supplementary Tables S1-S3.

PET data were acquired in list mode format. Following the microPET scan, a 10 min 80 kV/500 μA CT scan was performed for attenuation and scatter correction. The same microPET scan, was also used to estimate the blood radioactivity through a non-invasive image-derived input function (IDIF) to estimate the volume of distribution (*V*_T (IDIF)_) as a surrogate of *V*_T_.

Acquired PET data were reconstructed into 39 frames of increasing length (12×10s, 3×20s, 3×30s, 3×60s, 3×150s, and 15×300s). Images were reconstructed using a list-mode iterative reconstruction proprietary spatially variant resolution modeling in 8 iterations and 16 subsets of the 3D ordered subset expectation maximization (OSEM 3D) algorithm [25]. Normalization, dead time, and CT-based attenuation corrections were applied, and PET image frames were reconstructed on a 128×128×159 grid with 0.776×0.776×0.796 mm^3^ voxels.

### Image processing and analysis

Image processing and analysis were performed with PMOD 3.6 software (PMOD Technologies, Zurich, Switzerland) as previously described [11]. For spatial normalization of the PET/CT images, brain normalization of the CT image to a CT template was performed by adapting the previously described protocol [26]. We applied the volumes-of-interest (VOIs) described in previous studies [10, 11], adapted from the Waxholm atlas [27]. Using those VOIs, time activity curves (TACs) were then extracted for individual brain regions. The IDIFs were extracted from the PET images by generating a VOI in the lumen of the left ventricle at an early time frame exhibiting maximal activity in the heart and applying a threshold of 50% of the max signal as previously described [24, 28–30]. Standardized uptake value (SUV) was calculated as activity concentration [kBq/cc] divided by injected dose [kBq] per body weight [g].

These regional TACs and IDIFs were used to estimate *V*_T (IDIF)_ values through kinetic modeling. *V*_T (IDIF)_ was estimated by fitting the extracted TACs with standard two-tissue compartmental model (2TCM), with blood volume fraction (*V*_B_) fixed at 3.6% [10, 11] and Logan linear model to calculate *V*_T (IDIF)_. Pixelwise kinetic modeling was performed using the Logan graphical analysis in PXMOD (PMOD 4.2). The averaged parametric maps were subsequently overlaid onto a Waxholm MRI template.

### Tissue collection for Autoradiography

Animals were euthanized by decapitation while under anaesthesia. Brains were dissected and fresh-frozen in 2-metylbuthane at − 35°C for 2 min and preserved at −80°C. Coronal sections of 10 µm thickness were collected at 1.10 mm from the bregma according to Paxinos and Franklin [31] in triplicate using a cryostat (Leica, Germany) on Superfrost Plus slides (Thermo Fischer Scientific, USA).

### Autoradiography

[^3^H]CHDI-009R was provided by Novandi Chemistry AB (Sweden) with a A_m_ of 82 Ci/mmol. Sections were air-dried at room temperature, pre-incubated for 20 min with binding buffer (50mM Tris-HCl + 120mM NaCl + 5mM KCl + 2mM CaCl_2_ + 1mM MgCl_2_, pH 7,4) and air-dried using warm airflow. Dried sections were subsequently incubated with total binding (TB) solution (0.5 nM of [^3^H]CHDI-009R in binding buffer) and non-specific binding (NSB) solution (0.5 nM of [^3^H]CHDI-009R + 10 µM of cold ligand in binding buffer) for 1h at room temperature.

Subsequently, sections were washed three times for 10 min in ice-cold washing buffer (50mM Tris-HCl, pH 7,4), followed by a brief wash in ice-cold distilled water, and dried for 2h at room temperature. Finally, all sections were exposed on imaging plates (BAS-TR2025, Fujifilm, Japan) for 120 min. Radioactivity was detected with a phosphor imager (Fuji FLA-7000 image reader) and quantified based on intensity values calculated using commercial tritium standards (American Radiolabelled Chemicals Inc., USA). For all samples, the entire surface of the coronal brain slice was used for quantification. The specific binding (SB) was determined by subtracting NSB from TB, which were both measured in triplicate and subsequently averaged. The SB averages were used for statistical analysis. SB was set to 0 if the values of NSB and TB were below the limit of detectability of the calibration curve. Analysis was performed blind to conditions in Fiji, ImageJ (National Institute of Health, USA, version: 2.1).

### Statistical analysis

The Shapiro-Wilk test revealed that *in vivo* data were normally distributed, while *in vitro* data demonstrated a non-Gaussian distribution; therefore, non-parametric testing was performed on *in vitro* data. All data was visually inspected for outliers, followed, if necessary, by statistical outlier testing with ROUT (Q=0.5%). Two outliers in motor cortex for 9-month-old HET mice were identified and excluded. A Pearson’s correlation with simple linear regression was applied to *V*_T (IDIF)_ values from 2TCM and Logan kinetic modeling for model verification. A Spearman’s correlation with interpolation (sigmoidal, 4PL, X is concentration) was applied to injected masses and *V*_T (IDIF)_ values from Logan for visualization of a potential mass dose effect. Regular two-way ANOVA with *post hoc* Bonferroni correction for multiple comparisons was applied to analyze *V*_T (IDIF)_ value differences between genotypes.

The individual relative test-retest variability (rTRV) was calculated as follows:

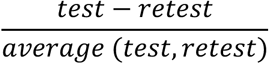

The individual absolute test-retest variability (aTRV) was calculated as follows:

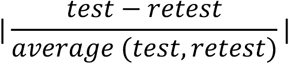

In addition, a Bland-Altman analysis was performed to assess the agreement between two measures on an individual level. A Mann-Whitney test was performed to investigate phenotypic differences in SB in autoradiography.

All statistical analyses above were performed with GraphPad Prism (version 9.2.0) statistical software. The Intraclass correlation coefficient (ICC) for the test-retest variability study was calculated with the software JMP Pro 17 (v17.0.0). Group and timepoint interaction was set as fixed effect. Animal ID, timepoint, interaction of animal ID and group, and interaction of animal ID and timepoint, were set as random effects. No effect was identified for either interaction. The ICC was reported as the ratio of the animal variance component for the effect to the sum of positive variance components. One-tailed power analyses were performed in G*Power (v3.1.9.6) with significance level (α) at 0.05, confidence (β) set to 80%. Data are represented as mean ± standard deviation (SD) or as mean ± standard error of the mean (SEM). All tests were two-tailed unless stated otherwise. Statistical significance was set at *p*<0.05 with the following standard abbreviations used to reference *p* values: ns, not significant; \**p*<0.05; \*\**p*<0.01; \*\*\**p*<0.001; \*\*\*\**p*<0.0001.

## Results

### [^11^C]CHDI-009R demonstrated high metabolic stability

[^11^C]CHDI-009R radiometabolite analysis revealed a high stability over time in both brain (e.g. 40 min: WT=90.9±8.6%; HET=93.8±2.0%; Fig. 2a) and plasma (e.g. 40 min: WT=95.1±1.7%; HET=96.4±1.2%; Fig. 2b) and with no observable peak other than the intact radioligand (Fig. S1). In addition, the plasma-to-whole-blood ratio was stable over time with no difference observed between genotypes (WT=1.8±0.1; HET=1.7±0.2; Fig. 2c). IDIF profiles of both genotypes displayed a comparable peak within 2 min of radioligand injection and a similar stable signal after 5 min p.i. until the end of acquisition (Fig. 2d). Given the high stability and no difference between genotypes, no corrections were applied to IDIFs for accurate *V*_T (IDIF)_ estimations.

**Fig. 2.**
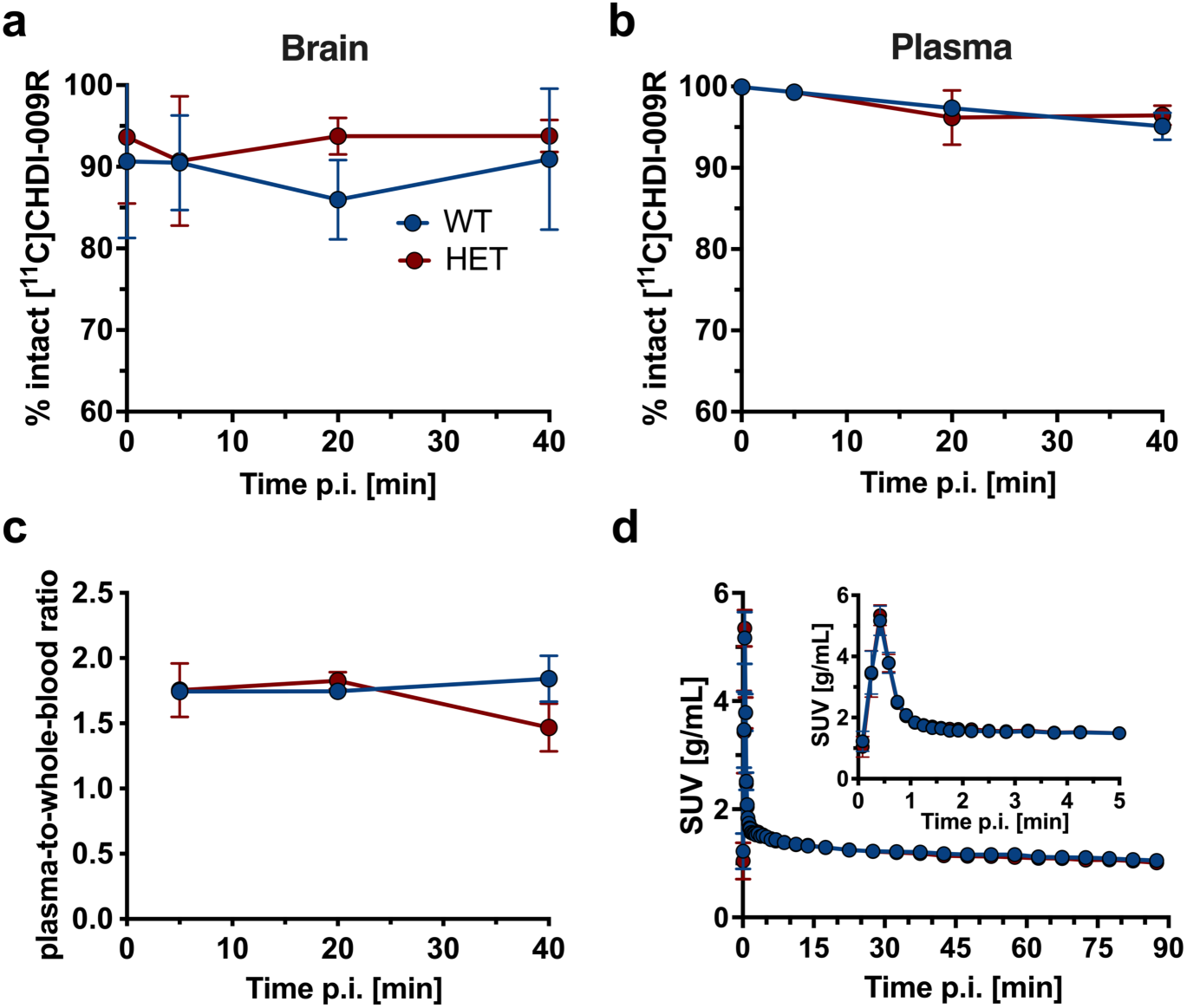
[^11^C]CHDI-009R brain and plasma analysis and IDIF of WT and HET zQ175DN mice. The intact radioligand in percentage for brain (a) and plasma (b) over time. c, Plasma-to-whole-blood ratio over time in both genotypes. d, Averaged IDIF SUV curves over 90 min scan time for both genotypes. d: Data are represented as mean±SEM. a-c: n=2-5 per genotype and timepoint; d: WT: n=20; HET: n=19

### [^11^C]CHDI-009R kinetic modeling with 90 min scan duration

The averaged striatal SUV TACs of [^11^C]CHDI-009R demonstrated rapid penetration into the brain in both genotypes, but a subsequent slower wash-out was observed in the brain in HET animals compared to WTs (Fig. 3a). We further observed an excellent agreement between 2TCM quantification and Logan graphical analysis (striatum: *r*^2^=0.996; *p*<0.0001) with minimal deviation from the identity line (slope = 0.998) (Fig. 3b); thus, supporting the use of Logan graphical analysis to quantify [^11^C]CHDI-009R in mice.

**Fig. 3.**
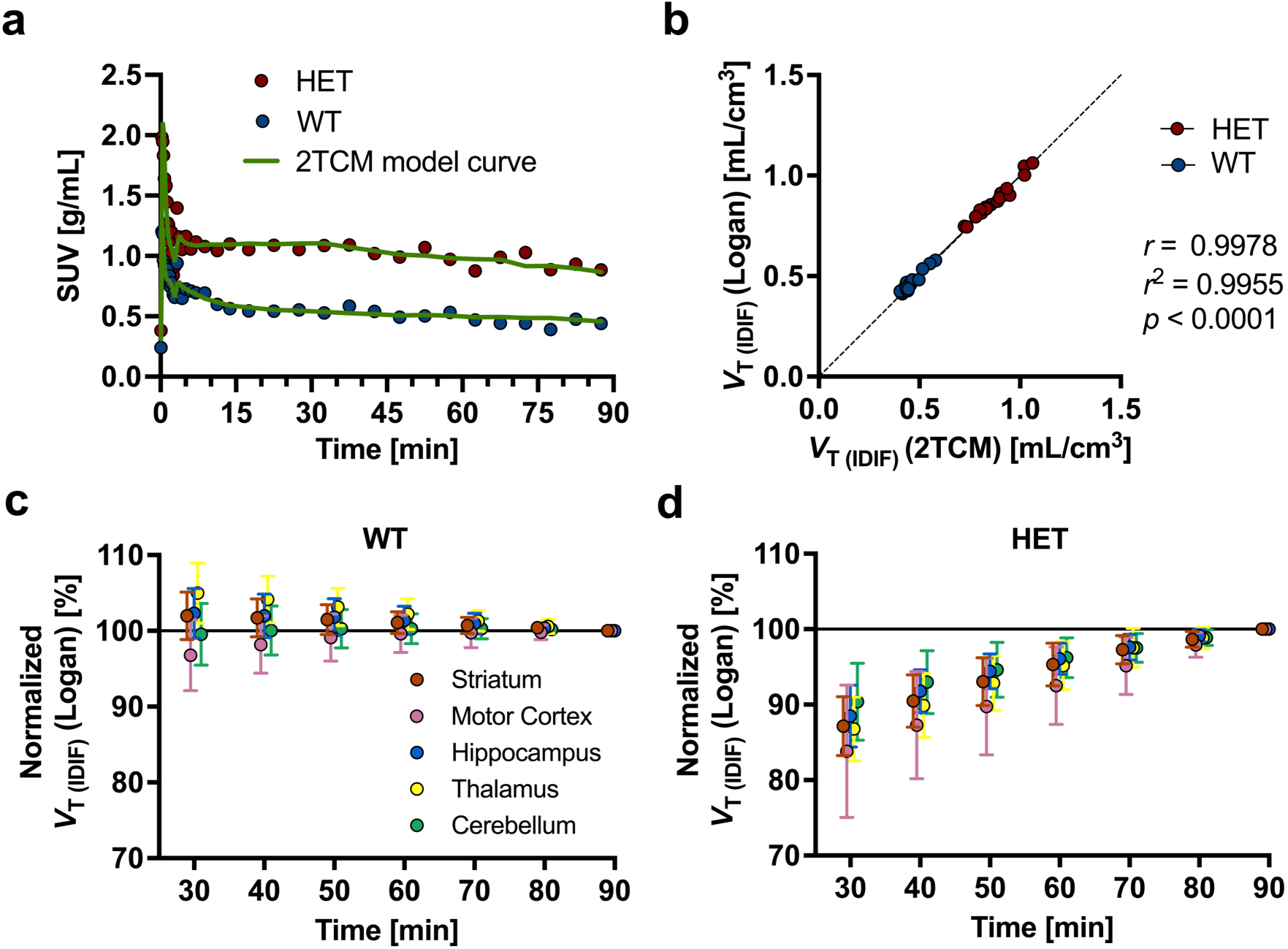
[^11^C]CHDI-009R brain kinetic modeling and V_T (IDIF)_ time stability with Logan. a, Exemplary striatal SUV TAC curves of WT and HET animals fitted to 2TCM. b, Pearson’s correlation of striatal V_T (IDIF)_ values with 2TCM and Logan for verification with simple linear regression. V_T (IDIF)_ time stability of Logan in WT (c) and HET (d) mice. V_T (IDIF)_ values were normalized to the 90 min scan acquisition time. b-d: WT: n=20; HET: n=19

Next, we performed a *V*_T (IDIF)_ time stability analysis to estimate the shortest recommended acquisition time. Although we observed no impact on *V*_T (IDIF)_ in WT animals (Fig. 3c), *V*_T (IDIF)_ in HET mice was susceptible to a steady underestimation, which passed the 90% threshold with a 60 min acquisition (Fig. 3d). Thus, shortening the scan acquisition time from 90 min was not recommended to avoid even minimal underestimation of a phenotypic effect. Further, we investigated the impact of injected mass on [^11^C]CHDI-009R *V*_T (IDIF)_. As shown in Fig. S2, a definite negative association was observed; thus, the recommended upper limit for injectable mass was set to 1.7 µg/kg.

### [^11^C]CHDI-009R quantification revealed phenotypic differences

The averaged parametric *V*_T (IDIF)_ maps of WT and HET mice shown in Fig. 4a visualized a clear genotype effect. The two-way ANOVA test revealed a significant genotype effect (F_(1, 183)_ = 987.3; *p*<0.0001) and the Bonferroni *post hoc* analysis demonstrated statistically significant increased signal in HET mice compared to their WT littermates in all tested brain regions (*p*<0.0001) (Fig. 4b). Phenotypic *V*_T (IDIF)_ differences ranged from 55.5±5.4% in the cerebellum to 105.8±6.2% in the thalamus (Table 1).

**Fig. 4.**
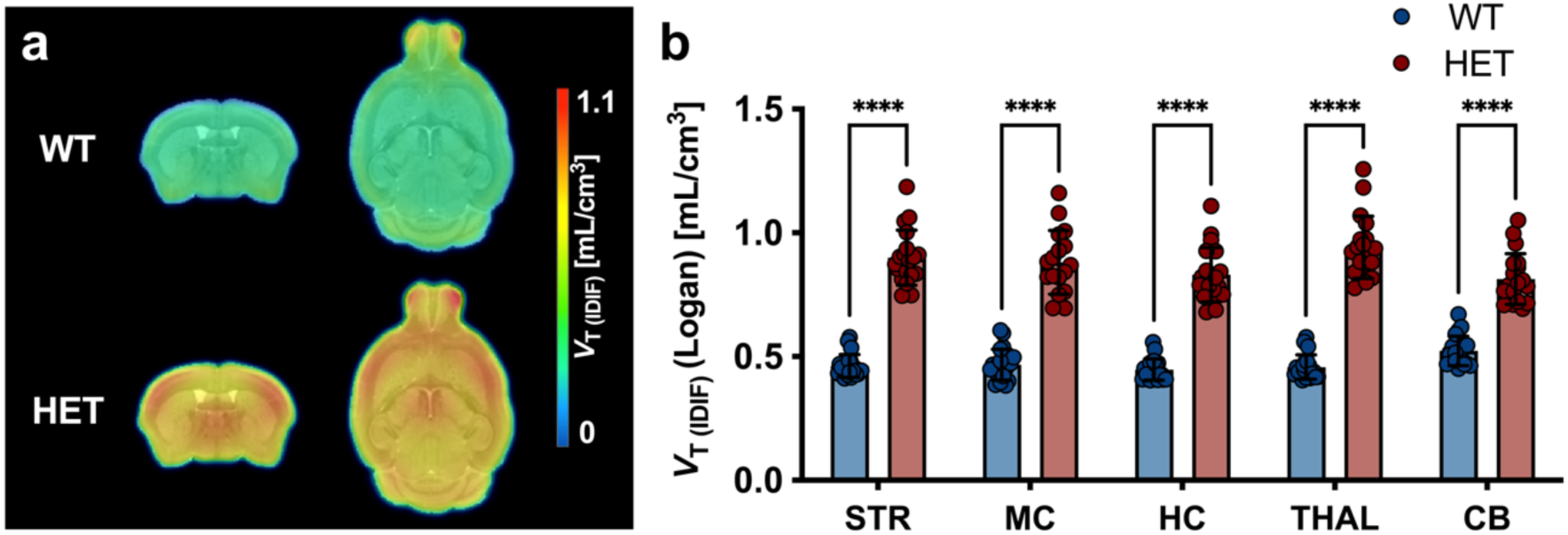
[^11^C]CHDI-009R mHTT aggregate quantification in 9-month-old mice. a, Group-averaged parametric maps for WT and HET zQ175DN mice overlaid onto MRI template. b, [^11^C]CHDI-009R V_T (IDIF)_ estimations in WT and HET mice. Two-way ANOVA with post hoc Bonferroni for multiple comparisons. ****p<0.0001. WT: n=20; HET: n=19. STR = Striatum; MC = Motor Cortex; HC = Hippocampus; THAL = Thalamus; CB = Cerebellum

**Table 1.** [^11^C]CHDI-009R V_T (IDIF)_ (Logan) [mL/cm^3^] quantitative values in 9-month-old mice.

| | WT<br>(mean±SD) | HET<br>(mean±SD) | Post hoc<br>*p* value | $\Delta\pm$SE [%] |
| --- | --- | --- | --- | --- |
| Striatum | 0.46±0.05 | 0.90±0.11 | <0.0001 | 94.6±6.2 |
| Motor Cortex | 0.47±0.06 | 0.88±0.13 | <0.0001 | 88.8±6.3 |
| Hippocampus | 0.45±0.04 | 0.83±0.11 | <0.0001 | 85.9±6.4 |
| Thalamus | 0.46±0.05 | 0.94±0.13 | <0.0001 | 105.8±6.2 |
| Cerebellum | 0.52±0.06 | 0.81±0.10 | <0.0001 | 55.5±5.4 |

WT: $n=20$ ; HET: $n=19$ . SD = standard deviation; $\Delta$ = phenotypic difference; SE = standard error

### [^11^C]CHDI-009R displayed excellent test-retest reproducibility

To measure the reliability of [^11^C]CHDI-009R, a test-retest study was performed in WT and HET zQ175DN mice at 9 months. Parametric *V*_T (IDIF)_ maps did not reveal any apparent differences between scans (Fig. 5a). The aTRV was overall low, indicating a high reliability, with a range from 5.1±4.0% to 7.7±4.2% in striatum of WT and HET animals (Fig. 5b; Table 2). The Bland-Altman plot displayed close-ranged levels of agreement (-16.0% to 14.7%) (Fig. 5c). In addition, the regional rTRV, with values ranging from -3.1±8.5% to 4.3±8.3% in WT and from -4.8±6.2% to 1.2±8.3% in HET, demonstrated limited difference across brain regions (Table 2). Overall, according to the ICC values, [^11^C]CHDI-009R displayed a good to excellent test-retest agreement (Table 2).

**Fig. 5.**
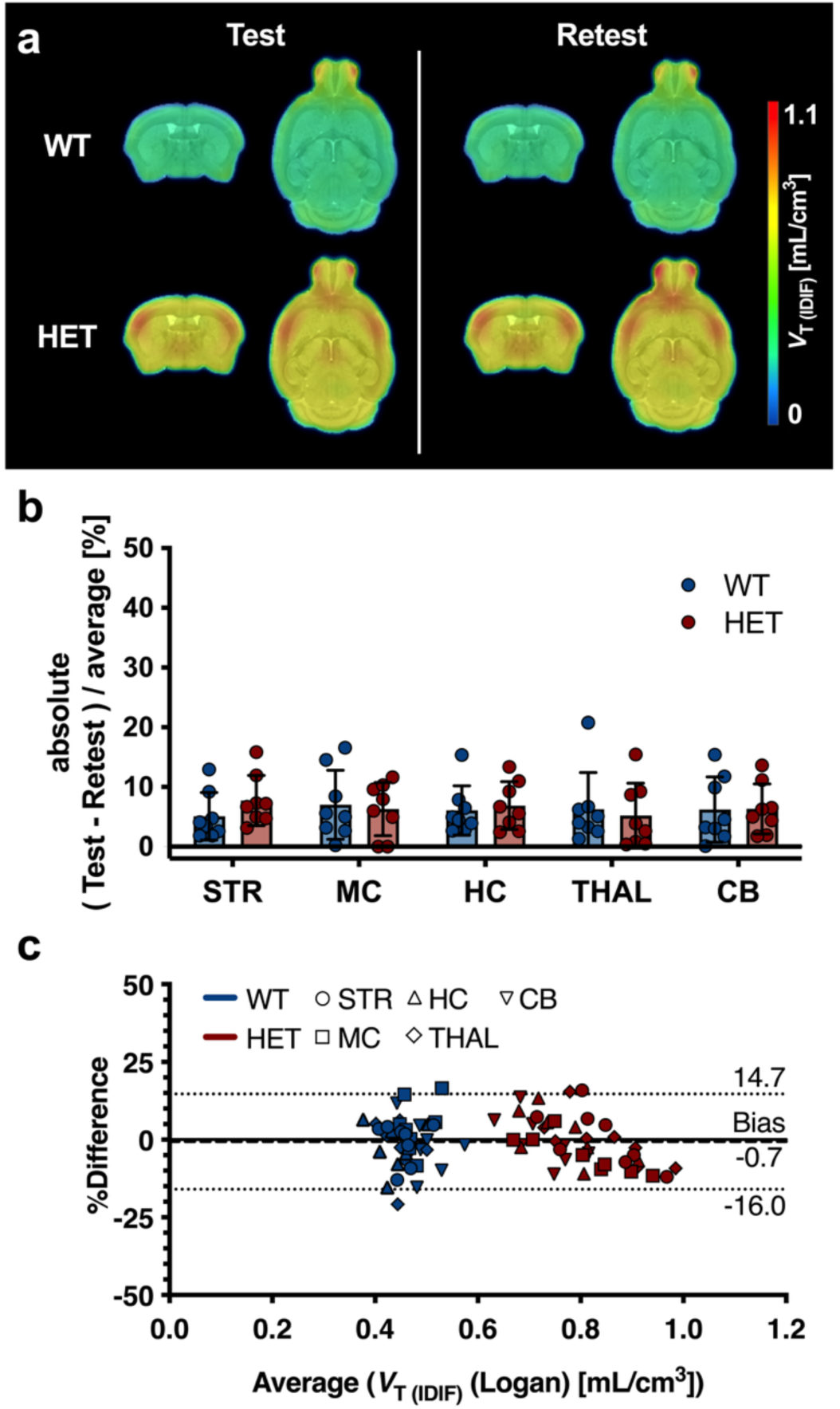
[^11^C]CHDI-009R test-retest variability analysis in 9-month-old mice. a, Group-averaged parametric maps for WT and HET zQ175DN mice at test and retest overlaid onto MRI template. b, Absolute test-retest variability analysis in percentage. c, Bland-Altman plot to display the level of agreement between test and retest. WT: n=8; HET: n=8. STR = Striatum; MC = Motor Cortex; HC = Hippocampus; THAL = Thalamus; CB = Cerebellum

**Table 2.** [^11^C]CHDI-009R test-retest V_T (IDIF)_ values and relative and absolute test-retest variability in 9-month-old mice.

| | Test $V_{T(IDIF)}$ (Logan)<br>[mL/cm<sup>3</sup>] | | Retest $V_{T(IDIF)}$ (Logan)<br>[mL/cm<sup>3</sup>] | | rTRV [%] | | aTRV [%] | | ICC |
| --- | --- | --- | --- | --- | --- | --- | --- | --- | --- |
|  | WT | HET | WT | HET | WT | HET | WT | HET |  |
| **Striatum** | 0.45±0.04 | 0.84±0.06 | 0.46±0.04 | 0.82±0.10 | -0.9±6.7 | 0.9±9.2 | 5.1±4.0 | 7.7±4.2 | 0.62 |
| **Motor Cortex** | 0.49±0.04 | 0.79±0.07 | 0.47±0.03 | 0.84±0.12 | 4.3±8.3 | -4.8±6.2 | 7.0±5.8 | 6.3± 4.5 | 0.79 |
| **Hippocampus** | 0.43±0.04 | 0.77±0.06 | 0.44±0.04 | 0.66±0.04 | -1.8± 7.4 | 1.2 ± 8.3 | 6.1± 4.2 | 6.9± 4.1 | 0.71 |
| **Thalamus** | 0.44±0.03 | 0.84±0.07 | 0.46±0.04 | 0.81±0.08 | -3.1±8.5 | -0.1±7.8 | 6.2± 6.2 | 5.2±5.4 | 0.72 |
| **Cerebellum** | 0.49±0.04 | 0.72±0.05 | 0.50±0.05 | 0.67±0.04 | -2.1± 8.3 | -0.1± 8.0 | 6.2± 5.5 | 6.3± 4.2 | 0.63 |

*rTRV = relative test-retest variability; aTRV = absolute test-retest variability; ICC = Intraclass correlation coefficient. Data is given as mean±SD. WT: n=8; HET: n=8*

### [^11^C]CHDI-009R differentiated genotypes at 3 months

In addition to the comprehensive assessment of [^11^C]CHDI-009R properties at 9 months, a study was performed at 3 months, when less mHTT aggregates are present in HET zQ175DN mice, providing insights into the degree of sensitivity for this radioligand. Based on parametric *V*_T (IDIF)_ maps, an increased signal in HET mice could be detected (Fig. 6a). Accordingly, the two-way ANOVA test revealed a significant genotype effect (F_(1, 175)_ = 62.3; *p*<0.0001). The Bonferroni *post hoc* analysis displayed statistically significant phenotypic differences between genotypes, ranging from 8.3±2.7% to 16.3±2.9% in test regions, with the exception of motor cortex (5.0±2.8%; not significant) (Fig. 6b; Table 3). The mHTT aggregate binding capacity of CHDI-009R was also demonstrated through a post-mortem autoradiography study with the tritiated version of the ligand. [^3^H]CHDI-009R displayed a significantly different SB in HET mice compared to WT littermates (Fig. 6c, *p*<0.0001).

**Fig. 6.**
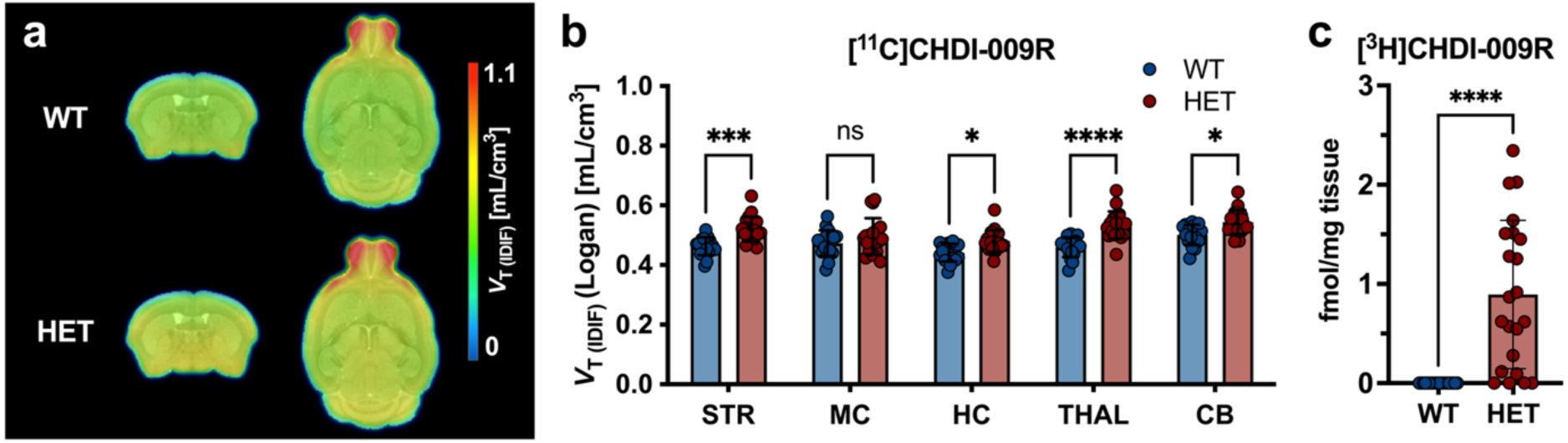
mHTT quantification in 3-month-old mice. a, Group-averaged parametric maps for WT and HET zQ175DN mice overlaid onto MRI template. b, [^11^C]CHDI-009R V_T (IDIF)_ estimations in WT and HET mice. c, [^3^H]CHDI-009R autoradiography of the whole brain in the same mice. ns=not significant; * p<0.05; *** p<0.001; **** p<0.0001. a and b: WT: n=18; HET: n=19; c: WT: n=20; HET: n=22. STR = Striatum; MC = Motor Cortex; HC = Hippocampus; THAL = Thalamus; CB = Cerebellum

**Table 3.** [^11^C]CHDI-009R V_T (IDIF)_ (Logan) [mL/cm^3^] quantitative values in 3-month-old mice.

| | WT<br>(mean±SD) | HET<br>(mean±SD) | Post hoc<br>*p* value | $\Delta\pm$SE [%] |
| --- | --- | --- | --- | --- |
| Striatum | 0.46±0.03 | 0.52±0.04 | 0.0002 | 12.3±2.9 |
| Motor Cortex | 0.47±0.04 | 0.50±0.06 | 0.3989 | 5.0±2.8 |
| Hippocampus | 0.44±0.03 | 0.48±0.04 | 0.0194 | 8.9±3.0 |
| Thalamus | 0.46±0.03 | 0.53±0.05 | <0.0001 | 16.3±2.9 |
| Cerebellum | 0.50±0.03 | 0.54±0.04 | 0.0104 | 8.3±2.7 |

WT: $n=18$ ; HET: $n=19$ . SD = standard deviation; $\Delta$ = phenotypic difference; SE = standard error

### [^11^C]CHDI-009R is suitable for evaluating therapeutic efficacy in zQ175DN mice

We performed power analyses to estimate group sizes needed for evaluating therapeutic efficacy (*α*=0.05, *β*=80%) in zQ175DN mice. As shown in Table S4, statistical power was good, as revealed by the range of Cohen’s *d* of 0.59 – 1.70 and 3.52 – 5.15 at 3 and 9 months, respectively. This indicated that at 9 months, [^11^C]CHDI-009R could significantly display a 20% difference in mHTT aggregates with acceptable group arms (e.g. striatum: *n*=21). At 3 months, however, only in striatum (*n*=23) and thalamus (*n*=26) could a larger effect (50% difference) between groups be detected with a similar sample size.

## Discussion

Here we report the *in vivo* characterization of a novel mHTT aggregate PET radioligand, [^11^C]CHDI-009R, with higher selectivity [20] than previously investigated radiotracers [12, 15, 16]. As in previous evaluation studies of mHTT aggregate radiotracers, we included only male mice for better comparison, as no sex differences have been observed in HET mice before [32]. Since the current study focused on evaluating the capability of [^11^C]CHDI-009R in detecting different levels of mHTT aggregates in the brain *in vivo*, only one sex was used. While research on sex differences is of significant importance, the inclusion of both sexes in this study was considered secondary.

We show that [^11^C]CHDI-009R rapidly enters the brain, followed by wash-out, indicating its reversible binding. The slower rate of radioligand wash-out from the brain of the HET animals was likely due to the presence of mHTT aggregates that are absent in WT. [^11^C]CHDI-009R demonstrated high stability in mouse plasma, and no metabolite correction was deemed necessary for IDIFs, although we could only obtain data until 40 min p.i. due to the loss of radioactive signal at later time points in the limited blood sample that can be collected as a consequence of the radioisotope’s half-life.

The cerebral retention time for both genotypes was suitable for estimating the *V*_T (IDIF)_, for which both 2TCM and Logan graphical analysis were shown as adequate models. The time stability kinetic analysis revealed that a reduction in scan acquisition time was not feasible without introducing a genotype-specific underestimation of the *V*_T (IDIF)_ values, which would result in an artificial decrease of phenotypic differences. The current scan duration of 90 min, however, borders the capability of obtaining data with limited noise due to the short half-life of the isotope (20.4 min), which requires a prompt scan acquisition and keeping the injected mass within the recommended limit of 1.7 µg/kg.

With those optimized PET imaging conditions, quantification with Logan indicated that [^11^C]CHDI-009R could significantly distinguish genotypes at both 9 and 3 months of age, with comparable magnitude as observed with previous mHTT radiotracers [10, 17]. Noteworthy, there was no change in *V*_T (IDIF)_ values of WT animals across brain regions, indicating that [^11^C]CHDI-009R can reliably and uniformly show the lack of a target in WT animals. Moreover, the post-mortem [^3^H]CHDI-009R autoradiography confirmed the lack of mHTT aggregates in WT mice, corroborating the previously described findings [20]. This represents an important indication of minimum off-target binding that could otherwise potentially diminish the utility of a radioligand, as shown for several Alzheimer’s disease-related radioligands [33].

Besides general affinity and selectivity, intra-subject variability was also an important factor to be considered. The reliability of [^11^C]CHDI-009R was demonstrated by the excellent test-retest performance as determined via Bland-Altman plot and ICC values [34]. This provides a significant benefit for a more accurate application, especially in longitudinal settings, and represents a favorable attribute compared to the poor reliability of [^18^F]CHDI-650 (ICC: 0.00 – 0.27) [17].

The capability of detecting early changes at the molecular level represents another important utility in the context of monitoring (early) therapeutic intervention, as described in the studies of HD mouse models [10, 32]. Our current work confirmed the *ex vivo* findings that [^3^H]-radiolabeled CHDI-009R [20] can detect statistically significant phenotypic differences at 3 months of age with both *in vivo* PET imaging and post-mortem autoradiography.

We have described several mHTT radiotracers that could identify a significant difference in striatal binding in 3-month-old zQ175DN HET mice [11, 17]. Especially [^11^C]CHDI-180R exhibited high metabolic stability and appropriate brain wash-out for imaging studies in both HD mouse models as well as in a large non-human primate (NHP) model of HD [35], suggesting [^11^C]CHDI-180R as a well-performing mHTT aggregate-directed PET radioligand for animal models [10, 14]. However, [^11^C]CHDI-180R demonstrated high intersubject variability and relatively low signal-to-background ratio in the first-in-patient PET study [19]. The increased SB and binding potential of [^3^H]CHDI-009R (SB=205.3±10.3 fmol/mg; B_max_=448 fmol/mg) compared to [^3^H]CHDI-180 (SB=70.9±11.0 fmol/mg; B_max_=163 fmol/mg) in HOM zQ175DN brains likely points towards higher sensitivity to detect mHTT aggregates with CHDI-009R, as shown in post-mortem HD brains [20]. Further, radio-binding assays revealed increased binding to mHTT aggregates with CHDI-009R (IC_50_=0.4±0.2 nM) in contrast to CHDI-180R (IC_50_=1.3±1.0 nM), while additionally minimizing the affinity for abeta- and/or tau-associated aggregates [20]. Thus, [^11^C]CHDI-009R demonstrates high potential for successful clinical translation given its improved characteristics over CHDI-180R. The utility of CHDI-009R as an imaging biomarker will be assessed in a clinical setting.

## Conclusions

[^11^C]CHDI-009R is a highly stable radioligand capable of detecting significant phenotypic differences between HET and WT zQ175DN mice at different ages in a reproducible manner. These findings suggest [^11^C]CHDI-009R to be a suitable radioligand to image mHTT aggregates in mice. Notably, a clinical investigation is underway to determine its suitability for assessing mHTT aggregates in humans.

## Declarations

### Ethics approval

Experiments followed the European Committee (decree 2010/63/CEE) and were approved by the Ethical Committee for Animal Testing (ECD 2019–39) at the University of Antwerp (Belgium).

### Availability of data and material

The PET datasets generated or analysed during the current study are available in the OpenNeuro repository, https://doi.org/10.18112/openneuro.ds006613.v1.0.1

## Acknowledgements

The authors thank Philippe Joye, Caroline Berghmans, Romy Raeymakers and Eleni Van der Hallen of the Molecular Imaging Center Antwerp (MICA) for their valuable assistance. FZ, LE, AVE, JA, AM, JV, SS, and DB are members of the µNeuro Research Centre of Excellence at the University of Antwerp. The authors acknowledge the contribution of the following CHDI staff: Drs Mette Skinbjerg, Robert Doot and Yuchuan Wang for helpful discussions; Drs Simon Noble and Vahri Beaumont for reviewing and providing constructive feedback; Brenda Lager for coordinating animal supply and Braden Boone for project management.

## Authors’ contributions

Conceptualization: JV, VK, JB, LL, SS, DB

Methodology: JV, AM, SS, DB

Data acquisition: FZ, LE, AVE, SDL

Formal analysis and investigation: FZ, LE, AVE

Data interpretation: FZ, JV, JA, VK, JB, LL, SS, DB

Writing – original draft preparation: FZ, SS, DB

Writing – review and editing: FZ, JV, SDL, JA, CD, VK, JB, LL, SS, DB Resources: CD, VK, JB, LL, SS, DB

Funding acquisition: SS, DB

Supervision: JV, SS, DB

## Competing interests

This work was funded by CHDI Foundation, Inc., a privately-funded nonprofit biomedical research organization exclusively dedicated to collaboratively developing therapeutics that improve the lives of those affected by Huntington’s disease. CD, VK, JB, and LL are employed by CHDI Management, Inc., the company that manages the scientific activities of CHDI Foundation. No other potential conflicts of interest relevant to this article exist.

## Funding

This work was funded by CHDI Foundation, Inc. under agreement number A-11627. DB acknowledges support by the Research Foundation Flanders (FWO) (ID: 1229721N). University of Antwerp supported the work through professorships for JV, SS, and DB.

## Supplemental material

**Fig. S1.**
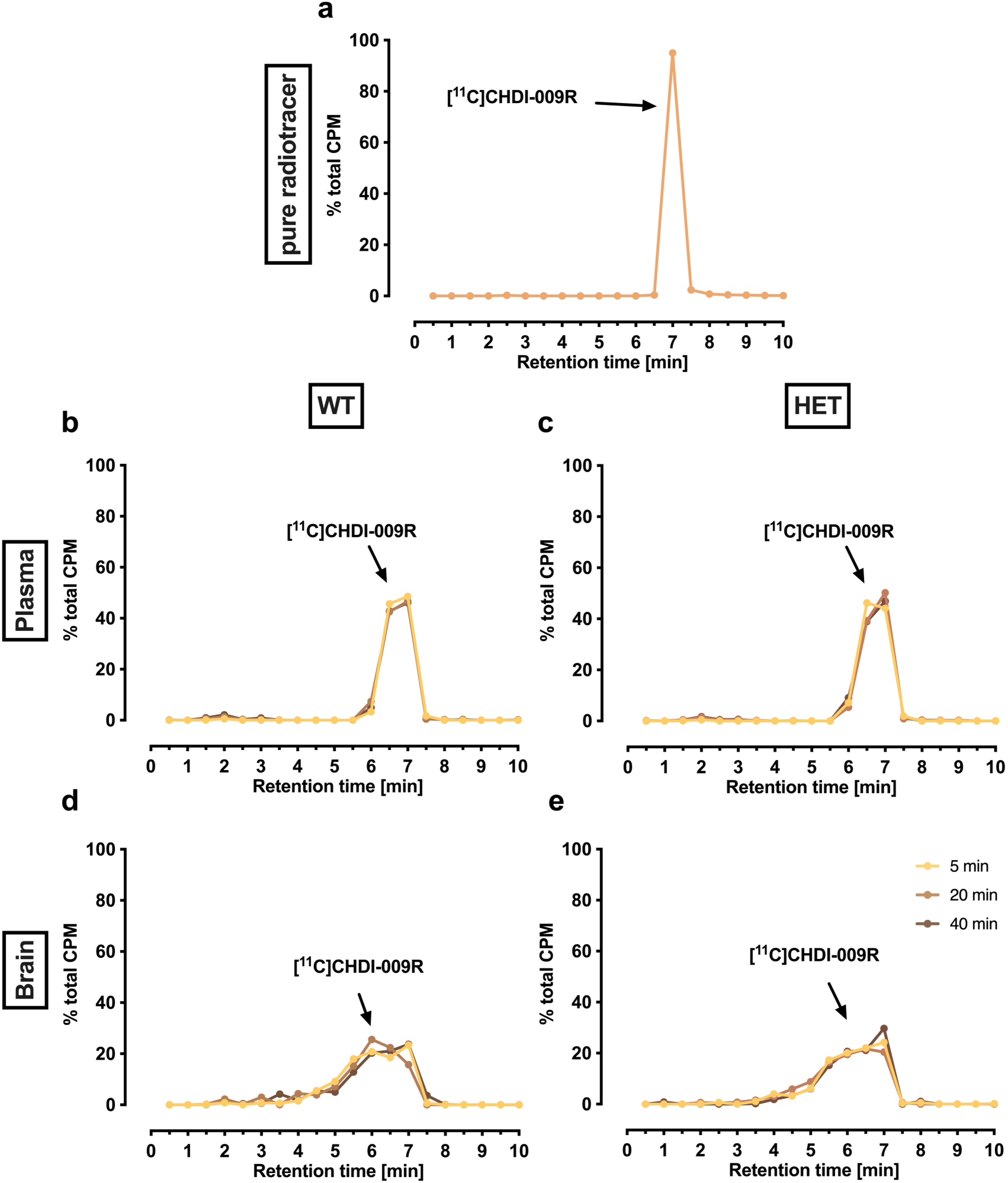
[^11^C]CHDI-009R radiometabolite analysis in WT and HET zQ175DN mice. a, Decay-corrected reconstructed radiochromatogram for the pure tracer. Decay-corrected reconstructed radiochromatograms for all time points in plasma (b and c) and brain (d and e) for WT (b and d) and HET mice (c and e). Data is given as the mean. n=2-3 per time point and genotype. CPM = counts per minute

**Fig. S2.**
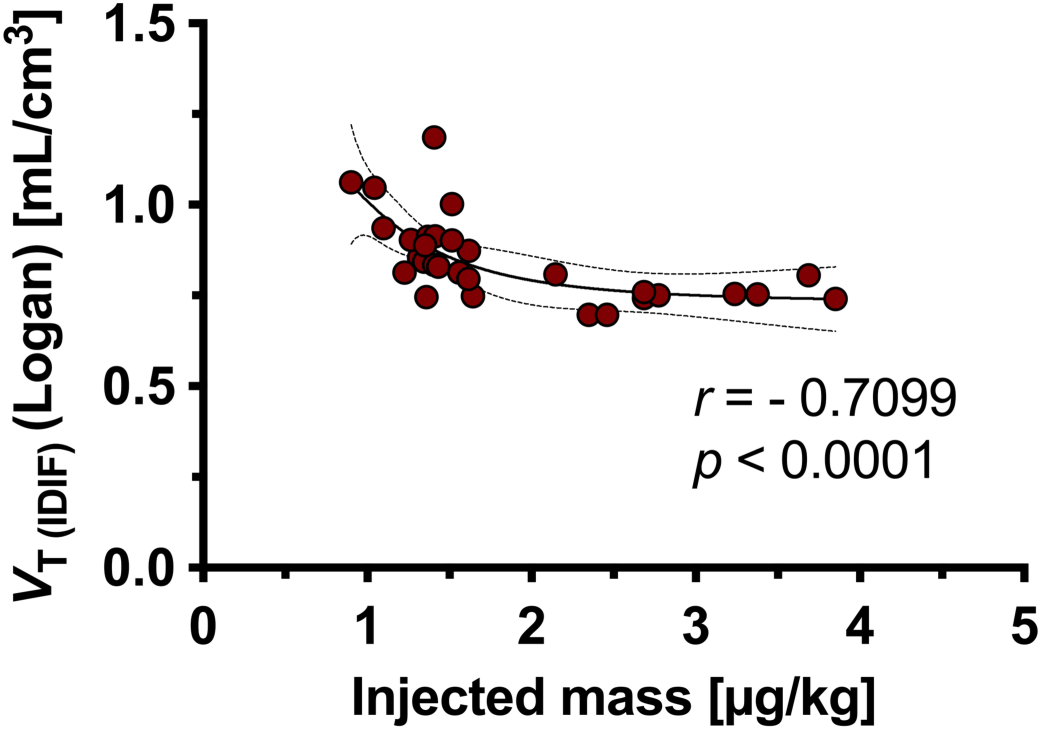
Mass dose effect in striatum of HET zQ175DN mice at 9 months. Spearman’s correlation of injected masses (µg/kg) and striatal Logan V_T (IDIF)_ values. The sigmoidal interpolation (sigmoidal, 4PL, X is concentration) estimates the influence of higher injected masses on the V_T (IDIF)_ value. The upper mass dose limit was recommended to be 1.7µg/kg. Data is represented as individual points. n=30

**Table S1.** Scan parameters during the PET imaging study in WT and HET zQ175DN mice for the test-retest study at 9 months.

|  | Test | Retest |  | Test | Retest |  |
| --- | --- | --- | --- | --- | --- | --- |
|  | WT (*n* = 8) |  | *p* value | HET (*n* = 8) |  | *p* value |
| Injected activity (MBq) | 6.3±1.6 | 7.1±2.5 | 0.1507 | 5.3±1.7 | 5.7±2.2 | 0.5148 |
| A<sub>m</sub> (GBq/μmol) at EOS | 112.6±28.6 | 122.8±26.9 | 0.3682 | 107.9±30.6 | 111.2±24.8 | 0.9375 |
| Weight (g) | 34.8±3.4 | 33.9±2.8 | 0.0006 | 30.8±2.6 | 30.1±2.2 | 0.0023 |
| Age (d) | 257.5±7.5 | 261.0±10.4 | <0.0001 | 261.1±9.5 | 267.0±13.9 | 0.0143 |
| Injected mass (μg/kg) | 1.31±0.15 | 1.27±0.17 | 0.6242 | 1.27±0.14 | 1.22±0.14 | 0.9275 |

Data is given as mean±SD. Unpaired t-test for comparison of genotypes. A<sub>m</sub> = molar activity; EOS = end of synthesis

**Table S2.** Animal and dose parameters for the PET imaging study in WT and HET zQ175DN mice for the evaluation at 9 months.

|  | WT (n = 20) | HET (n = 19) | p value |
| --- | --- | --- | --- |
| Injected activity (MBq) | 3.8±0.6 | 4.5 ± 1.2 | 0.0062 |
| A<sub>m</sub> (GBq/μmol) at EOS | 71.2±16.1 | 73.2 ± 16.8 | 0.7122 |
| Weight (g) | 29.7±1.8 | 34.8 ± 2.8 | <0.0001 |
| Age (d) | 263.3±4.3 | 263.2 ± 2.7 | 0.9221 |
| Injected mass (μg/kg) | 1.38 ± 0.20 | 1.35 ± 0.28 | 0.9337 |

Data is given as mean±SD unless stated otherwise. Unpaired t-test for comparison of genotypes.
A<sub>m</sub> = molar activity; EOS = end of synthesis

**Table S3.** Animal and dose parameters for the PET imaging study in WT and HET zQ175DN mice for the evaluation at 3 months.

|  | WT (n = 18) | HET (n = 19) | p value |
| --- | --- | --- | --- |
| Injected activity (MBq) | 5.0±0.9 | 4.8±0.9 | 0.4608 |
| A<sub>m</sub> (GBq/μmol) at EOS | 100.8±16.6 | 99.1±17.9 | 0.7676 |
| Weight (g) | 28.4±2.2 | 27.7±1.8 | 0.2975 |
| Age (d) | 108.0±3.3 | 106.6±4.1 | 0.2523 |
| Injected mass (μg/kg) | 1.31 ± 0.14 | 1.34 ± 0.19 | 0.6164 |

Data is given as mean±SD. Unpaired t-test for comparison of genotypes. A<sub>m</sub> = molar activity;
EOS = end of synthesis

**Table S4.** [^11^C]CHDI-009R power analysis of 3- and 9-month-old mice.

| age [M] | Brain region | striatal $\Delta \pm SE$ [%] | Cohen's $d$ | 10% | 20% | 30% | 40% | 50% | 100% |
| --- | --- | --- | --- | --- | --- | --- | --- | --- | --- |
| **3** | **Striatum** | 12.3±2.9 | 1.70 | 551 | 139 | 62 | 36 | 23 | 6 |
| | **Motor Cortex** | 5.0±2.8 | 0.59 | 4947 | 1238 | 540 | 310 | 199 | 37 |
| | **Hippocampus** | 8.9±3.0 | 1.13 | 1238 | 310 | 136 | 78 | 51 | 11 |
| | **Thalamus** | 16.3±2.9 | 1.70 | 632 | 159 | 70 | 41 | 26 | 6 |
| | **Cerebellum** | 8.3±2.7 | 1.13 | 1238 | 310 | 136 | 78 | 51 | 11 |
| **9** | **Striatum** | 95.8±6.1 | 5.15 | 78 | 21 | 10 | 6 | 4 | 2 |
| | **Motor Cortex** | 88.8±6.3 | 4.05 | 125 | 32 | 15 | 9 | 6 | 2 |
| | **Hippocampus** | 85.9±6.4 | 4.59 | 105 | 27 | 13 | 8 | 5 | 2 |
| | **Thalamus** | 105.8±6.2 | 4.87 | 92 | 24 | 11 | 7 | 5 | 2 |
| | **Cerebellum** | 55.5±5.4 | 3.52 | 148 | 38 | 17 | 10 | 7 | 3 |

*One-tailed power analysis with $\alpha=0.05$ and $\beta=80\%$ . Animal numbers needed for significance*
*between treated and non-treated HET groups in a mHTT-lowering study. 100% equals difference*
*between non-treated HET and WT groups. 3 months: WT: $n=18$ ; HET: $n=19$ ; 9 months: WT:*
*$n=20$ ; HET: $n=19$ . M = months; $\Delta$ = phenotypic difference; SE = standard error*

## Notes

### Summary of Updates

References updated due to recent publications: Delva, A. et al. PET imaging with [(1)(1)C]CHDI-00485180-R, designed as radioligand for aggregated mutant huntingtin, in people with Huntington's disease. Eur J Nucl Med Mol Imaging, 2025. Liu, L., et al. Isoindolinone-Based PET Tracers for Imaging Mutant Huntingtin Aggregates. J Med Chem, 2025. 68(14): p. 14818-14842. Certain graphs moved from the supplemental file to the main text.

